# Expression Analysis, Molecular Characterization and Prognostic Evaluation on *TMED4* and *TMED9* Gene Expression in Glioma

**DOI:** 10.1101/2022.04.07.487570

**Authors:** Md. Asad Ullah, Tahani Tabassum, Maisha Farzana, Abu Tayab Moin, Umme Salma Zohora, Mohammad Shahedur Rahman

## Abstract

Here, we utilized a database mining approach to unfold the prognostic and therapeutic potentials of Transmembrane EmP24 Trafficking Protein 4 (*TMED4*) and 9 (*TMED)* coding gene expressions in glioma. Both the genes were found to be overexpressed at the mRNA and protein level in low grade glioma (LGG) and glioblastoma multiforme (GBM) tissues including different glioma cell lines. Significant increase in the expression level of these genes with advancing glioma patients’ age, glioma grades and histological subtypes was observed. Differential and distinct promoter and coding sequence methylation pattern of *TMED4* and *TMED9* was observed in LGG and GBM tissues that may aid in methylation-sensitive diagnosis of glioma patients. The presence of multiple heterozygous genetic alterations (frequency: 0.4-1.1%) in those genes unveiled their potentials in high-throughput screening of glioma patients. The overexpression of *TMED4* and *TMED9* genes was associated with poor overall survival (OS) of LGG and GBM patients (HR:>1.6). Association of the expression levels of these genes with different immune cell infiltration levels i.e., B cell and T cell and modulators like CD274 and IL10RB was observed providing assurance in *TMED*-based diagnostic measure and therapeutic intervention discovery. Furthermore, functional enrichment analysis of the neighbor genes of *TMED4* and *TMED9* revealed that they are involved in metal ion binding, focal adhesion of cells and protein processing, and the deregulation of these activities are associated with gliomagenesis. Altogether, this study suggests that *TMED4* and *TMED9* are potential prognostic and therapeutic targets for glioma. However, further laboratory research is warranted.

## 1. Introduction

Brain tumor, the commonest solid tumor in children, refers to a diverse group of neoplasms emerging from distinct anatomic sites of the intracranial tissues and the meninges with varying degrees of malignancy ranging from benign to aggressive [1]. Malignant and non-malignant brain tumors are rare comprising only 2% of all cancers but are amongst the most fatal cancers accounting for substantial mortality and morbidity across the world [2]. Glioma, a broad category of diffusely infiltrative brain and spinal cord tumor arising from the glial cells, is the most common primary malignant brain tumor in adults with a very poor prognosis. About 33% of all brain tumors are gliomas, out of which approximately 70%-80% are malignant [3,4]. Global incidence of glioma varies greatly, displaying a higher incidence in men than women, and a higher occurrence across the Western population compared to the Asian or African population [5].

In accordance with the presumed cell of origin and histological features, gliomas can be broadly classified into astrocytomas including glioblastoma, oligodendrogliomas, ependymomas, and mixed gliomas [6-8]. Amongst these, glioblastoma multiforme (GBM), also commonly referred to as glioblastoma, accounts for approximately 60% to 70% of malignant gliomas and is the most frequently occurring primary astrocytomas [9]. World Health Organization (WHO) has categorized gliomas based on their histopathological characteristics such as cytological atypia, anaplasia, mitotic activity, microvascular proliferation, and necrosis [3]. According to this standard for defining the nomenclature, diagnosis, and malignancy of glioma, GBM is the most aggressive, invasive glioma subtype and by definition a high-grade tumor (HGG, WHO grade IV) [3,10]. Although the global incidence of GBM is less than 10 per 100,000 people, it’s extremely poor prognosis with a median survival time period of 14-15 months approximate from diagnosis makes it a serious health threat [11]. Despite several international therapeutics approaches, GBM treatment is still the most challenging task in clinical oncology with very limited success in the range of different treatments investigated [12]. Low-grade glioma (LGG, WHO grade I-IV), brain tumors arising from two different cell types namely astrocytes and oligodendrocytes, are another diverse group of primary brain tumors that often arise in young otherwise healthy people and is the slowest growing glioma types in adults [3]. LGGs account for 6.4% of all the primary CNS tumors, and although it rarely causes major neurological deficits besides seizures, recent data suggests that LGG can grow at a continuous rate reaching 4 to 5 mm per year [13,14]. Continuous growth of this untreated symptomatic LGG can eventually undergo malignant transformation triggering a more complicated disease course, reduced quality of life, and worse prognosis [14], raising serious complications and health concerns. Although LGG and GBM remains the most devastating form of brain cancer, the proper management and treatment of glioma patients is often interrupted by a lack of proper understanding of glioma pathogenesis and the protective structure of the CNS, as well as the unavailability of efficient diagnostic and therapeutic measures [15,16]. Therefore, there is an ever-increasing demand to discover an effective molecular target for appropriate and accurate stratification and therapeutic interventions in glioma patients.

The Transmembrane EmP24 Domain-Containing Proteins (TMED), also known as p24 proteins, are members of a family of sorting receptors abundantly present in the cellular subcompartments of the early secretory pathway including the endoplasmic reticulum (ER), Golgi body, and the intermediate compartment of all representatives of the domain Eukarya [15-17]. This family of proteins has an average molecular mass of 24 kDa and are central regulators in cell signaling, the secretion of biomolecules, and intracellular protein transport [17]. Due to their active influence in determining the composition, structure, and function of the ER and the Golgi, as well as their role in maintaining cellular traffic, they have been associated with a variety of human diseases, including carcinomas. For example, *TMED3* promotes hepatocellular carcinoma cell proliferation in by regulating STAT-signaling pathway [18, 19]. *TMED2* is associated with poor prognosis of liver hepatocellular carcinoma, lung adenocarcinoma and head-neck squamous cell carcinoma. Again, TMED3 overexpression is associated with poor clinical outcome in lung squamous cell carcinoma and colorectal cancer patients’ survival [20-22]. Considering the significant role of TMED proteins in cellular proliferation and differentiation, all of the ten known TMED proteins have been studied in tumour-associated contexts [23-25], and have displayed a promising role as biomarkers of different cancer types. Amongst these proteins, *TMED4* and *TMED9* have been less investigated for their role in brain tumor pathomechanism. However, a recent study screened TMED9 proteins for unleashing their link to vascular invasion and poor prognosis with hepatocellular carcinoma, displaying promising results [26]. Moreover, the prognostic potential of TMED9 protein has been investigated in another recent study demonstrating an association between high expression of *TMED9* and breast carcinoma cell proliferation and migration [29]. Another study reported significant upregulation of TMED4 proteins in longer-surviving patients with pancreatic ductal adenocarcinoma (PDAC) [30].

In this study, we have investigated the differential expression of *TMED4* and *TMED9* genes within different subgroups of glioma patients and associated different molecular aspects of these proteins with the glioma patients’ clinical outcomes. We have aimed to unleash the role of *TMED4* and *TMED9* genes and their post-transcriptional products as prognostic markers and therapeutic targets in brain cancer gliomas, more specifically LGGs and GBM. The scientific findings of this study should assist in TMED-based therapeutic and diagnostic measure formulation in glioma.

## 2. Materials and Methods

### 2.1. Differential Expression Analysis on *TMED4* and *TMED9* Genes in LGG and GBM Tissues

The mRNA level expression pattern of *TMED4* and *TMED9* genes in LGG and GBM tissues was determined using two servers i.e., GEPIA 2 (http://gepia2.cancer-pku.cn/) and OncoDB server (http://oncodb.org/) to increase the fidelity of the findings. GEPIA 2 is an online platform that allows the RNA sequencing data analysis on cancerous and normal tissues from integrated The Cancer Genome Atlas (TCGA) and Genotype-Tissue Expression (GTEx) samples [31]. OncoDB also assists in differential expression analysis of different genes in cancerous tissues and correlating gene expression with cancer patient’s clinical outcome [32]. The expression pattern of the genes in different glioma cell line was analyzed using the Expression Atlas server (https://www.ebi.ac.uk/gxa/home) [33]. Finally, the protein level expression of *TMED4* and *TMED9* genes was determined from the Human Protein Atlas (HPA) (https://www.proteinatlas.org/) server by exploring the tissue and pathology modules containing the immunohistochemistry images of gene expression [34].

### 2.2. Expression Analysis of *TMED4* and *TMED9* Genes in Accordance with Glioma Patients’ Demographic and Clinical Features

The expression level of our genes of interest in relation to glioma patients’ age, glioma grades (WHO), Isocitrate Dehydrogenase (IDH) mutation status and histological subtypes of glioma was determined using the online platform Chinese Glioma Genome Atlas (CGGA) (http://www.cgga.org.cn/). This is a web-based user-friendly database enabling users to compare the expression of mRNA, DNA methylation and miRNA levels between normal and brain tumor samples [35]. This server also allows the analysis of correlation between different gene/mRNA level features with brain cancer patients’ clinical manifestation by performing Analysis of Variance (ANOVA) test in between the control and test cohorts. The result was considered significant based on the p value cutoff of <0.5. Finally, the co-expression pattern of *TMED4* and *TMED9* in glioma tissues was also evaluated from this server.

### 2.3. Analysis of the Promoter and Coding Sequence Methylation Pattern in *TMED4* and *TMED9* Genes in LGG and GBM Tissues

The genomic profiles of *TMED4* and *TMED9* in LGG and GBM tissues delineating their methylation status was retrieved from the OncoDB server. The result displayed significant methylation position across the coding sequence of the selected genes. Thereafter, the methylation pattern of *TMED4* and *TMED9* gene was compared to the sample type and histological type of TCGA LGG (n=530) and TCGA GBM samples (n=631), respectively from the UCSC Xena Browser. UCSC Xena browser in an online data repository that allows the users to explore genomics data and compare genotypic and phenotype variables [36]. Finally, the association between *TMED4* and *TMED9* methylation and glioma patients’ survival rate was discovered utilizing the Gene Set Cancer Analysis (GSCA) web-based tool (http://bioinfo.life.hust.edu.cn/GSCA). GSCA is an integrated tool that helps the users in identifying correlation between specific gene expression and genomic variation, patients’ clinical features, immune infiltration level and so on [37]. The results obtained from this served were considered significant based on a p-value cutoff of >0.05.

### 2.4. Analysis of the Frequency of Mutation and Copy Number Alterations in *TMED4* and *TMED9* Genes in Glioma Tissues

The mutation and Copy Number Alteration (CNA) frequency in *TMED4* and *TMED9* in LGG and GBM tissues was identified from the cBioPortal Server (https://www.cbioportal.org/) [38]. In the server, the genomic data deposited for LGG and GBM by TCGA, CPTAC, MSK, UCSF and a few other organizations spanning a total of 7 studies incorporating 2,404 samples were selected for the analysis. Thereafter, the OncoPrint and cancer types summary were investigated to observe the mutation and CNA events in those selected genes in glioma tissues. Finally, the association of CAN events in *TMED4* and *TMED9* genes with the glioma patients’ Overall Survival (OS), Disease Specific Survival (DSS), Relapse-free Survival (RFS) and Progression-free Survival (PFS) was evaluated from the GSCA server. The correlation in this step was analyzed based on hazard ratio and considered significant for p<0.05.

### 2.5. Survival Analysis on Glioma Patients in Relation to *TMED4* and *TMED9* Expression

Initially, the OS of LGG and GBM patients in accordance with *TMED4* and *TMED9* expression was compared in the PreCog using the individual cohort of this server (https://precog.stanford.edu/) [39]. This server enables the users to assess genomic data and find out correlations with cancer patients’ clinical sequalae. In other words, this server allows to define whether upregulation or downregulation of a particular gene is favorable or unfavorable for cancer patients’ OS and DSS. To further increase the credibility of survival analysis, the association between the expression level of our gene of interest and glioma patients’ OS was established from the OncoLnc server (http://www.oncolnc.org/) that helps linking the survival information on cancer patients with mRNA, miRNA and lncRNA expression levels from TCGA cohorts [40]. In either case, the survival data were analyzed based on HR and considered significant based on p values cutoff of >0.05.

### 2.6. Analysis on the Association between Immune Infiltration Level and *TMED4* and *TMED9* Expressions in Glioma Patients

Initially, the association between the selected gene expression and different immune cell infiltration levels i.e., B cell, T cell, Macrophage, Natural Killer (NK) cells in glioma tissues was determined from the immune module in TIMER 2 web-based tool (http://timer.cistrome.org/). TIMER 2 is a publicly available database that helps users in identifying specific gene expression with immune cell abundance levels based on multiple immune deconvolution strategies [41]. In the next step, the correlation between *TMED4* and *TMED9* expression and different immune modulator expression in LGG and GBM tissues was identified from the TISIDB online portal (http://cis.hku.hk/TISIDB/). This platform allows the interaction analysis between tumor and immune system b enabling the users to explore the association between different gene expression immunomodulators expression like chemokine, immune-receptors, immune-inhibitors [42]. In both cases, the results were analyzed based on spearman correlation coefficient and considered significant on the basis of p value.

### 2.7. Examining the Co-expressed Neighbor Genes of *TMED4* and *TMED9* in Glioma Tissues and Their Functional Enrichment Analysis

The co-expressed neighbor genes of *TMED4* and *TMED9* in glioma tissues was evaluated from the LinkedOmics server (http://linkedomics.org/login.php) using the TCGA LGG and TCGA GBM cohorts [43]. Initially, top positively correlated genes were inspected on the basis of pearson correlation coefficient. Thentop 10 highest positively co-expressed genes were selected from each cohort and utilized in the subsequent functional enrichment analysis using the Enrichr server [44]. Eventually, the co-expression mode of the top selected genes in glioma tissues was inspected using the CGGA server.

## 3. Results

### 3.1. mRNA and Protein Level Expression of *TMED4* and *TMED9* in LGG and GBM

The selected genes were investigated to understand their expression level at the mRNA and protein levels. The GEPIA 2.0 analyses revealed that *TMED4*mRNA is highly expressed both in LGG tissues (Tumor:518, Normal:207) and GBM tissues (Tumor:163, Normal:207) than their respective normal counterparts (**Figure 1a**). The similar inspection for *TMED9* gene also revealed that its transcriptional product is highly expressed both in LGG and GBM tissues compared to the normal brain tissues (**Figure 1b**). Thereafter, the *TMED4* and *TMED9* genes were also analyzed in the OncoDB database to observe their expression patterns in two forms of cancerous samples. Similar to the previous step, it was observed that *TMED4* mRNA is highly expressed in LGG tissues (p=3.31e-21) and GBM (p=2.1e-30) tissues when compared to the normal samples (**Figure 1c** and **1d**). For T*MED9*, LGG tissues (p=3.9e-129) and GBM tissues (1.3e-72) also showed higher expression in comparison with normal tissues (**Figure 1e** and **1f**). Interestingly, most noticeable differences between LGG and GBM cohorts were observed for *TMED9* expression along with the highest fold change in GBM tissues as observed from the results obtained from both servers. Afterward, the expression pattern of the genes was analyzed across different glioma cell lines. As in par with the previous results, *TMED9* showed higher expression in most of cell lines than *TMED4* (**Figure 1f)**. Finally, the protein level expression of the *TMED4* and *TMED9* genes was inspected by analyzing the immunohistochemistry (IHC) images from the HPA server. Herein, the normal central nervous system (CNS0 tissues showed low staining for the IHC antibody administered against the *TMED4* and *TMED9* proteins, whereas, cancerous tissues showed medium to high IHC intensity reflecting their higher protein level expression (**Figure 2)**.

**Figure 1:**
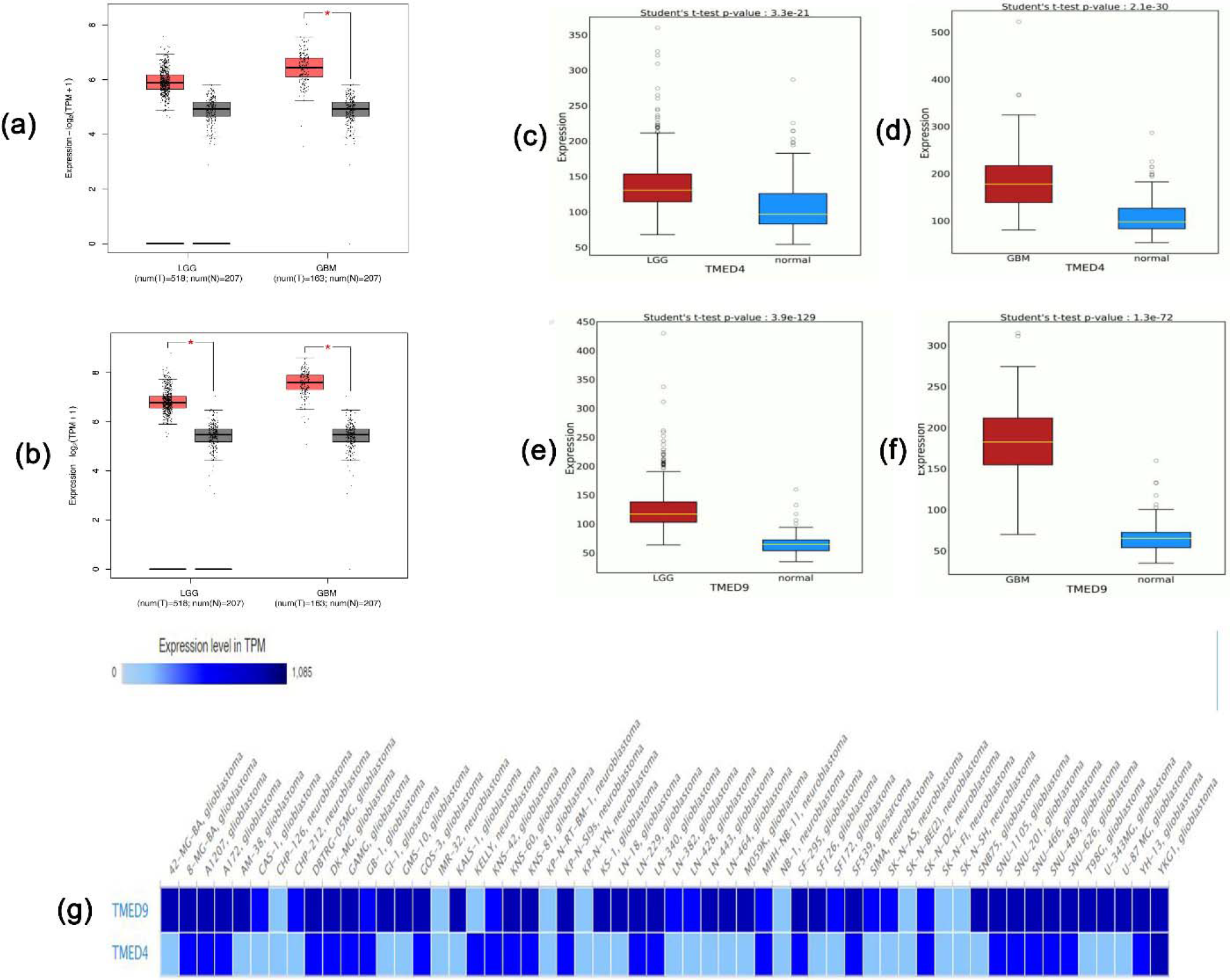
The mRNA level differential expression of *TMED4* (a) and *TMED9* (b) in LGG and GBM from GEPIA 2.0 server. Red and black boxes representing cancerous and normal samples, respectively (results are normalized with Log2 transformation). The OncoDB comparison of *TMED4* expression in LGG (c) and GBM (d) tissues and *TMED9* expression in LGG (e) and GBM (f) tissues in contrast to normal CNS tissues (results are presented in transcript per million (TPM) unit). The expression pattern of *TMED4* and *TMED9* in different glioma cell lines(results are presented in TPMunit) (g).

**Figure 2:**
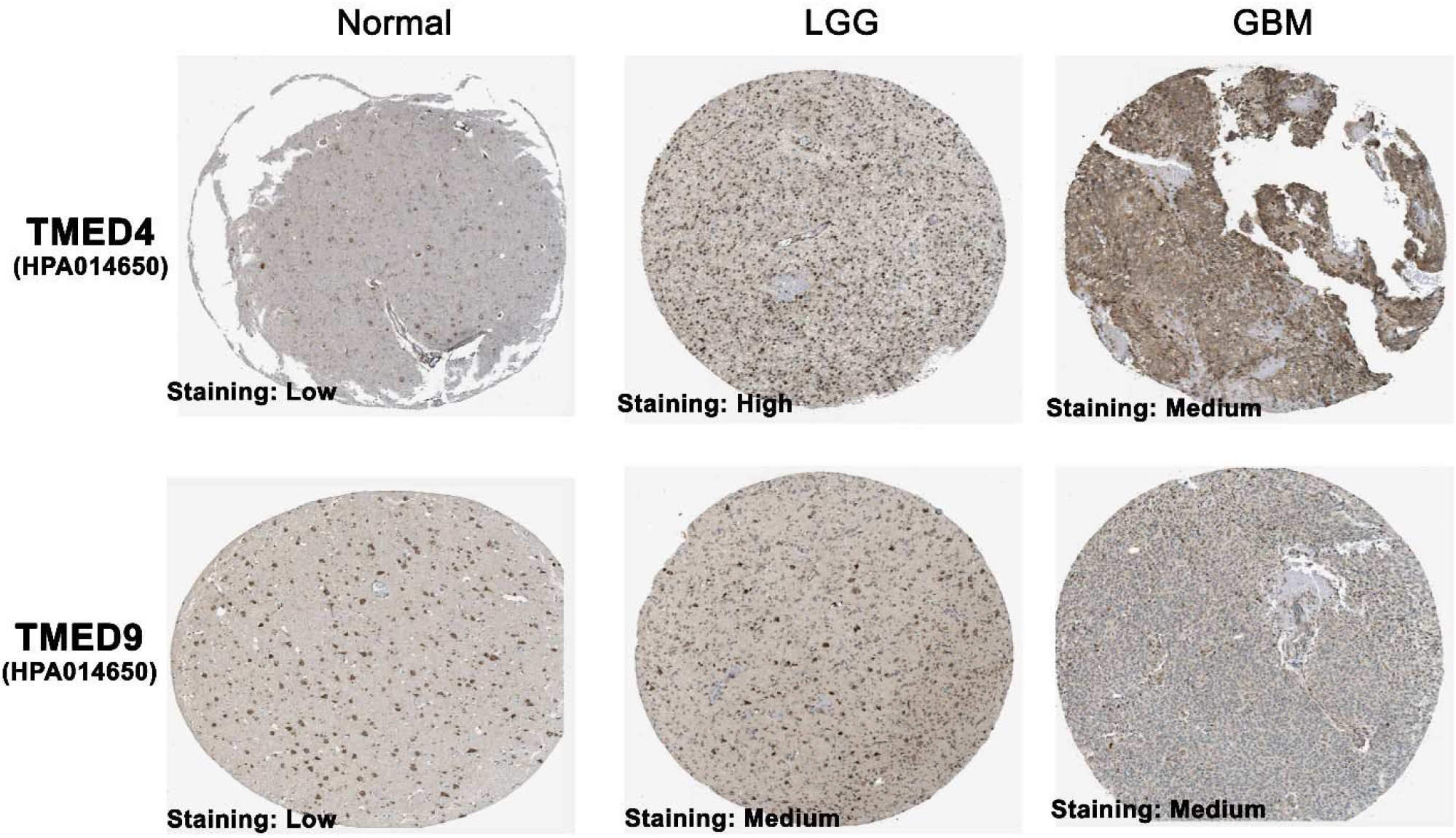
The post-mortem IHC images representing protein level expression of *TMED4* and *TMED9* in LGG, GBM and adjacent normal tissues. Normal tissues showed lower protein expression whereas LGG and GBM samples showed higher expression.

### 3.2. Expression Pattern of *TMED4* and *TMED9* across Different Grades, Subtypes and Demographic Conditions of Glioma Patients

In this step, the *TMED4* and *TMED9* genes were enquired in the CGGA server to discover their expression across different clinical and demographic conditions of glioma patients. Significant association between the expression levels and patients’ age was observed (**Figure 3**). Both the *TMED4* (p=2.5e-07) and *TMED9* (p-1.2e-06) were found be expressed at a higher level in >42 years age group compared to the <42 years group. Moreover, upward trend in *TMED4* (p=1.3e-11) and *TMED9* (p=1.9e-26) expression was also observed across World Health Organization (WHO) grade II, III and IV gliomas (**Figure 3**). Considering the prevalence of Isocitrate Dehydrogenase (IDH) mutation in glioma patients, the expression level of the selected genes in this study was also analyzed in relation to IDH mutation status. Interestingly, wildtype glioma tissues showed significantly high level of expression of both *TMED4* (p=5.9e-11) and *TMED9* (p=2.8e-22) genes (**Figure 3**). Thereafter, their differential expression pattern across different histological subtypes was determined. Similar pattern of increment in those gene expression in accordance with advancing subtype was observed. In both the cases, *TMED4* (p=1.1e-08) and *TMED9* (p=1.1e-08) showed least expression in Oligoastrocytoma tissues followed by a gradual rise in expression level with a few events of downward trends and finally achieving maximum expression level in Glioblastoma (**Figure 3**). Finally, the co-expression pattern of *TMED4* and *TMED9* was analyzed and it was found that both the genes are highly co-expressed in primary (Cor: 0.609, p: 2.65e-24) and recurrent glioma (Cor: 0.577, p: 9.36e-07) tissues (**Supplementary Figure S1**).

**Figure 3:**
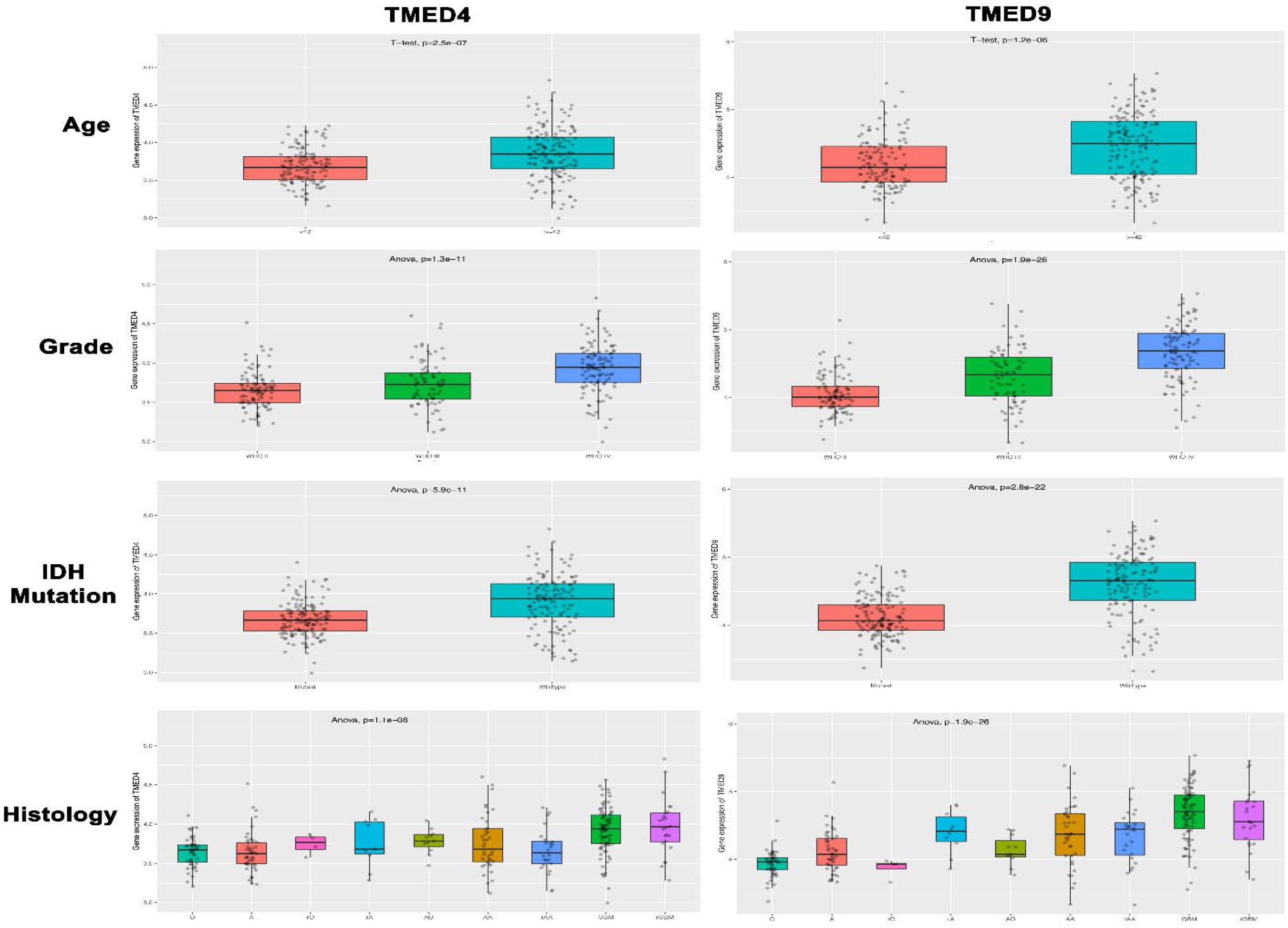
The expression pattern of *TMED4* and *TMED9* genes across glioma patients’ age, glioma grades, IDH mutation status and histological subtypes. Significant association between the expression of the genes and all selected items was observed. IDH: Isocitrate dehydrogenase, O: Oligoastrocytoma, A: Astrocytoma, rO: Recurrent Oligoastrocytoma, rA: Recurrent Astrocytoma, AO: Anaplastic Oligoastrocytoma, AA: Anaplastic Astrocytoma, rAA: Recurrent Anaplastic Astrocytoma, GBM: Glioblastoma, rGBM: Recurrent Glioblastoma.

### 3.3. The Promoter and Coding Sequence Methylation Pattern of *TMED4* and *TMED9* Genes in LGG and GBM Tissues

The promoter and coding sequence methylation status of *TMED4* and *TMED9* genes was retrieved from the OncoDB server. *TMED4* gene’s promoter region was found to be less methylated in LGG tissues whereas the 3’ end of the coding region showed highest methylationlevel (**Figure 4a**). Similar pattern of methylation was also observed for *TMED4* in GBM tissues, however, a delicate difference both in the promoter region and downstream landscape to promoter was observed (**Figure 4b**). In case of *TMED9*, a completely different pattern of methylation was observed when compared to *TMED4* gene methylation. The result suggested that the promoter region of *TMED9* is less methylated than *TMED4* in LGG tissues (**Figure 4c**). However, unlikely to *TMED4*, no observable difference in methylation pattern of *TMED9* in GBM tissues in contrast to the LGG tissues was observed (**Figure 4d**). Thereafter, their coding sequence methylation pattern was analyzed from the UCSC Xena browser. The analysis result from this server supported the findings of previous steps by revealing that most of CpG islands might be present at the 3’ end of the coding sequence as observed in the red colored landscape (**Supplementary Figure S2**). The result also indicated that both the genes have distinct methylation pattern which might be maintained through the transition between LGG and GBM states. Finally, the relation between *TMED4* and *TMED9* methylation and glioma patients’ survival rate was observed. Significant association between *TMED9* hypomethylation and GBM patients’ poor overall survival (OS) (p=0.0046) and disease specific survival (DSS) (p=0.0095) was observed (**Figure 4e** and **4f**).

**Figure 4:**
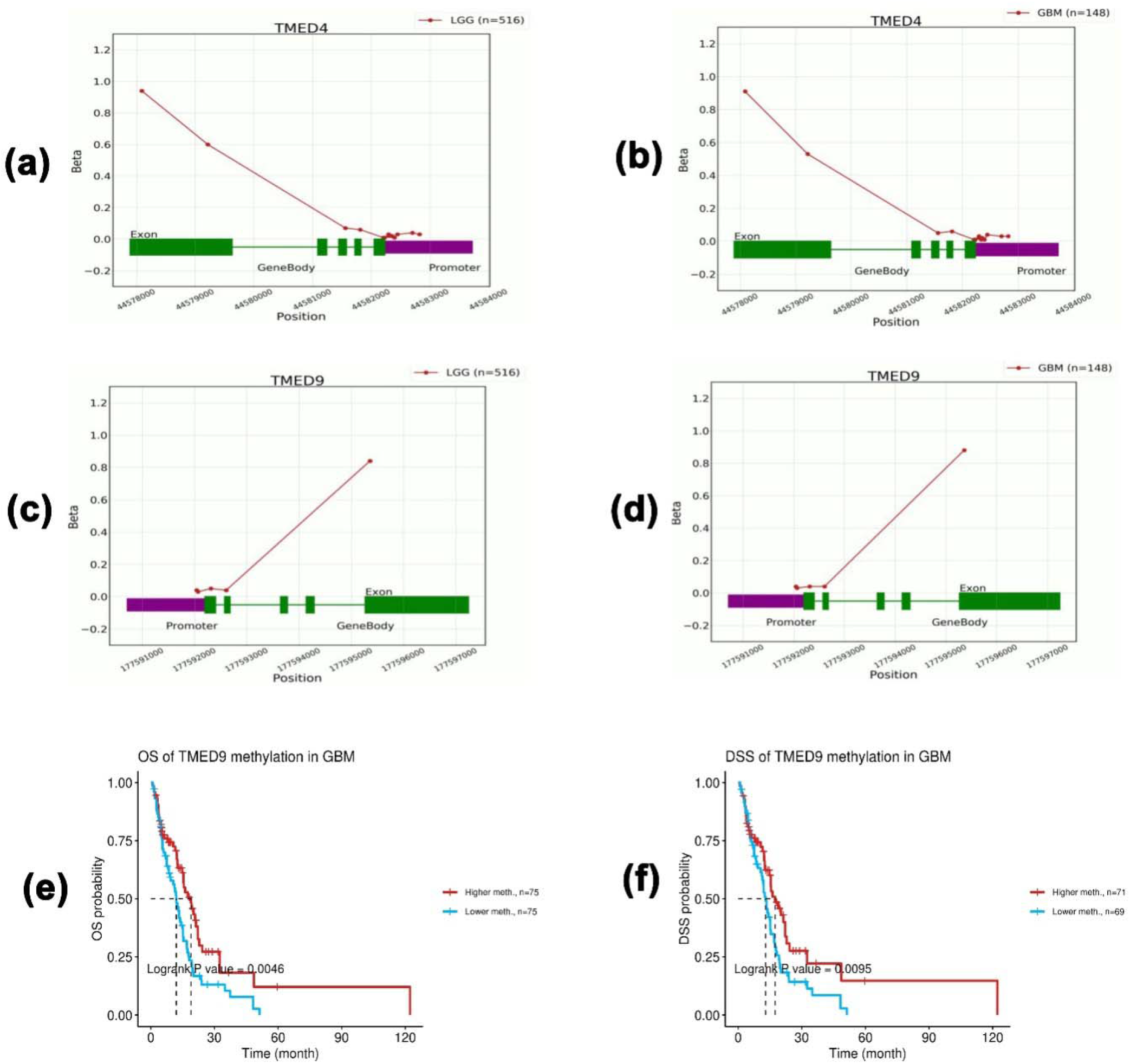
The result of methylation pattern analysis of *TMED4* gene in LGG (a) and GBM tissues (b) and *TMED9* gene in LGG (c) and GBM (d) tissues. Distinct pattern of promoter and coding sequence methylation was observed for both the genes in the selected tissue types. The association between *TMED9* mutation and GMB patients’ overall survival (e) and disease specific survival (f).

### 3.4. The Frequency of Mutation and Copy Number Alterations in *TMED4* and *TMED9* Genes in LGG and GBM Tissues

The mutation and copy number alteration (CNA) frequency in *TMED4* and *TMED9* genes in glioma tissues was determined using the cBioPortal server. Overall, *TMED4* (frequency:1.1%) was found to undergo more CNA events than the *TMED9* (frequency: 0.4%) gene in glioma tissues (**Figure 5a**). Moreover, amplification was the only CNA event in *TMED4*, whereas, *TMED9* experienced deep deletion along with amplification in different selected glioma studies (**Figure 5a and 5b**). However, no synonymous or nonsynonymous mutation was recorded for the selected genes in glioma tissues.Thereafter, the type of amplification and deletion in *TMED4* and *TMED9* genes in glioma tissues was determined from the GSCA server. It was found that, *TMED4* experienced more amplification events in the GBM tissues compared to the LGG tissues. Furthermore, majority of the amplification was heterozygous while homozygous amplification contributed to a minor portion (**Figure 5c**). On the contrary, *TMED9* gene underwent more heterozygous deletion than amplification both in the LGG and GBM tissues. Finally, the association between CNA in those genes and glioma patients’ survival was evaluated from the GSCA server. Although *TMED9* did not show any significant correlation, *TMED4* CNA was discovered to be negatively associated with both the LGG and GBM patients’ OS, DSS and progression free survival (PFS) (p<0.001) (**Figure 5d**) (**Supplementary Table S1**).

**Figure 5:**
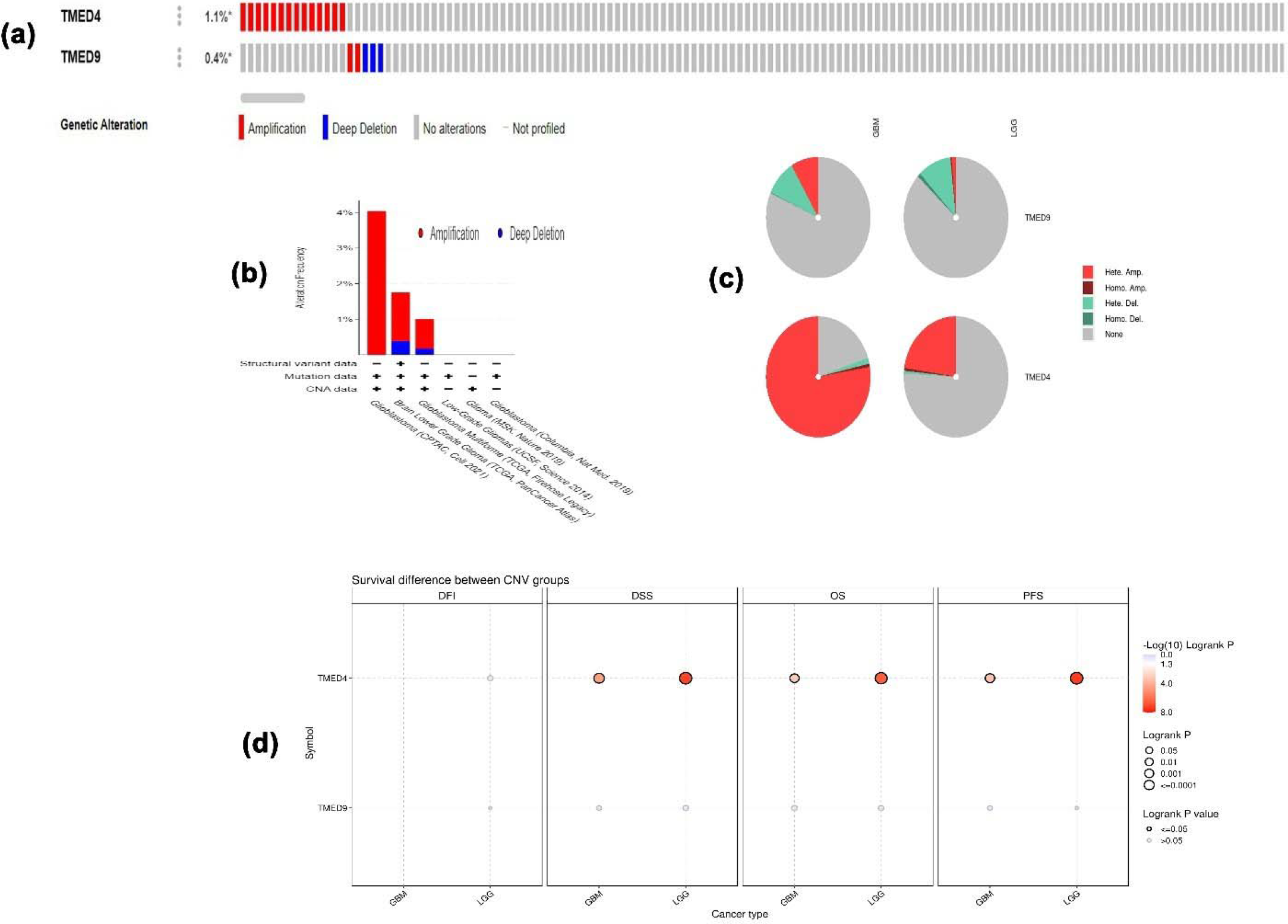
The oncoprint summary of the CNA events in *TMED4* and *TMED9* genes in glioma tissues (a). Type of CNA events found in *TMED4* and *TMED9* genes in glioma tissues (b). Classification of individual CNA events observed in the selected genes (c). The association between the CNA events in *TMED4* and *TMED9* genes as glioma patients’ different survival rate (d).

### 3.5. Association between *TMED4* and *TMED9* Expression and Glioma Patients’ Survival Rate

In this step, the *TMED4* and *TMED9* genes were explored to examine the association between their expression and LGG and GBM patients’ OS rate in the PreCog server. *TMED4* gene expression was found to negatively affect the OS of LGG patients (Hazard Ratio (HR): 4.59, p<0.0001) (**Figure 6a**). Similar to the LGG patients, *TMED4* expression was also discovered to be responsible for poor OS of GBM patients (HR: 1.12) (**Figure 6b**). However, the association was not found to be statistically significant as evidenced by higher p value of 0.12. In case of *TMED9* in LGG patients, high expression was again significantly accounted for the worse OS of the patients (HR: 3.47, p<0.0001) (**Figure 6c**). Finally, high expression of *TMED9* was found to be negatively correlated with the OS of GMB patients too (HR: 1.86, p=0.048) (**Figure 6d**). To further validate the findings of survival analysis from PreCog server we carried out the OS analysis again with the help of OncoLnc server. As in par with the previous findings, *TMED4* overexpression was revealed to be the significant cause of poor prognosis in LGG (p=4.55e-07) and GBM patients (p=0.003) (**Supplementary Figure S3a** and **S3b**). Similarly, significant negative association of *TMED9* overexpression in LGG (p=3.85e-06) and GBM patients’ (p=0.018) OS was also reported (**Supplementary Figure S3c** and **S3d**).

**Figure 6:**
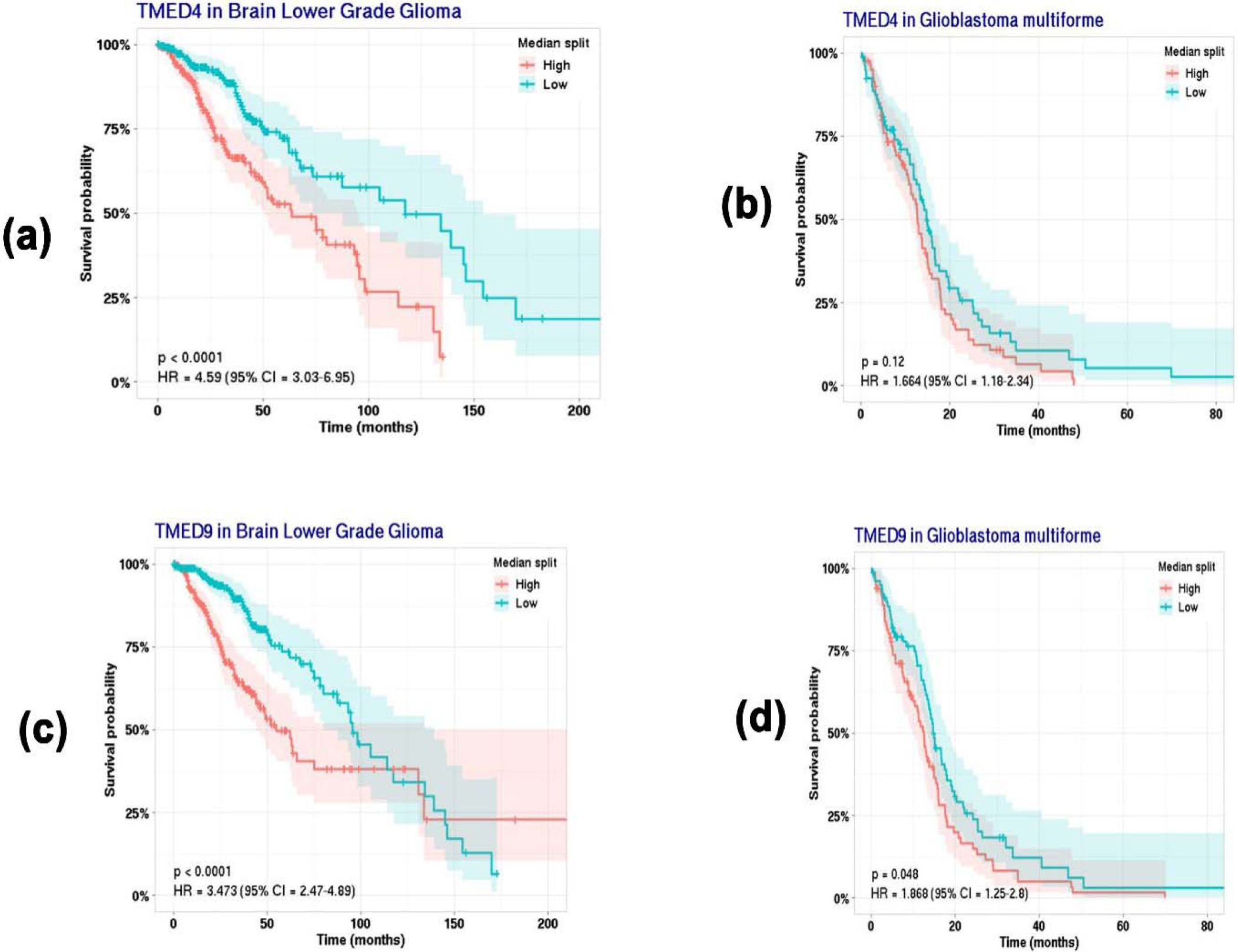
The result of overall survival analysis on *TMED4* gene expression in LGG(a) and GBM (b) patients from PreCog server. *TMED4* overexpression was found to be negatively correlated with poor OS of LGG and GBM patients (p<0.05). The overall survival analysis on *TMED9* gene expression in LGG (c) and GBM (d) patients which revealed negative correlation.

### 3.6. Association between *TMED4* and *TMED9* Expression and the Abundance of Tumor-infiltrating Immune Cells in LGG and GBM Patients

The association between the selected gene expression and tumor-infiltrating immune cell abundance was identified from the TIMER 2 server. Significant positive correlation between *TMED4* expression and B cell infiltration was observed in LGG patients (Cor:0.09, p=3.20e-02) (**Figure 7a**). Moreover, *TMED4* expression was significantly and positively associated with the CD8+ T cell (Cor:0.13, p=4.21e-03) infiltration level in LGG patients, whereas, a negative correlation was identified for Natural Killer (NK) cell (Cor: -0.389, p=1.07e-18). In GBM patients, *TMED4* did not show any significant positive or negative correlation with any of the immune cell infiltration level (**Figure 7b**). On the contrary, *TMED9* expression was negatively correlated with the B cell (Cor: -0.39, p=4.27e-19) and CD8+ T cell (Cor: 0.44, p=1.78-24) abundance in LGG patients (**Figure 7c**). Significant positive correlation between *TMED9* expression and CD4+ T cell (Cor: -0.39, p=4.27e-19), Macrophage (Cor: -0.39, p=4.27e-19) and NK cell (Cor: -0.39, p=4.27e-19) abundance was observed in LGG patients (**Figure 7c**). Interestingly, *TMED9* expression was negatively correlated with B cell (Cor: -0.27, p=1.42e-03), CD8+ T cell (Cor: -0.38, p=4.77e-06), CD4+ T cell (Cor: -0.12, p=1.51e-02), and NK cell (Cor: -0.26, p=2.18-103) infiltration level in GBM patients (**Figure 7c**). Moreover, significant positive correlation for *TMED9* expression was recorded only for Macrophage abundance in GBM patients (Cor: 0.20, p=1.48e-02). Finally, the expression level of *TMED4* and *TMED9* was explored to identify the association with different immunoinhibitor i.e., IL10RB, CD274 expression in glioma cells from TISIDB server. Significant association between *TMED4* expression and IL10RB (Cor: 0.121, p=0.005) and CD274 (Cor: 0.157, p=3e-03) abundance in LGG tissues was observed (**Figure 8a** and **8b**). IL10RB expression also showed positive correlation with *TMED4* gene expression in GBM tissues (Cor: 0.548, p<2.2e-16) (**Figure 8c)**. For *TMED9* expression, significant association with IL10RB (Cor: 0.204, p=0.008) and CD274 (Cor: 0.161, p=0.038) expression in LGG tissues was reported (**Figure 8e** and **8f**). Additionally, IL10RB (Cor: 0.454, p=1.17e-09) and CD274 (Cor: 0.298, p=1e-02) expression in GBM tissues were significantly and positively correlated with *TMED9* expression levels (**Figure 8g** and **8h**).

**Figure 7:**
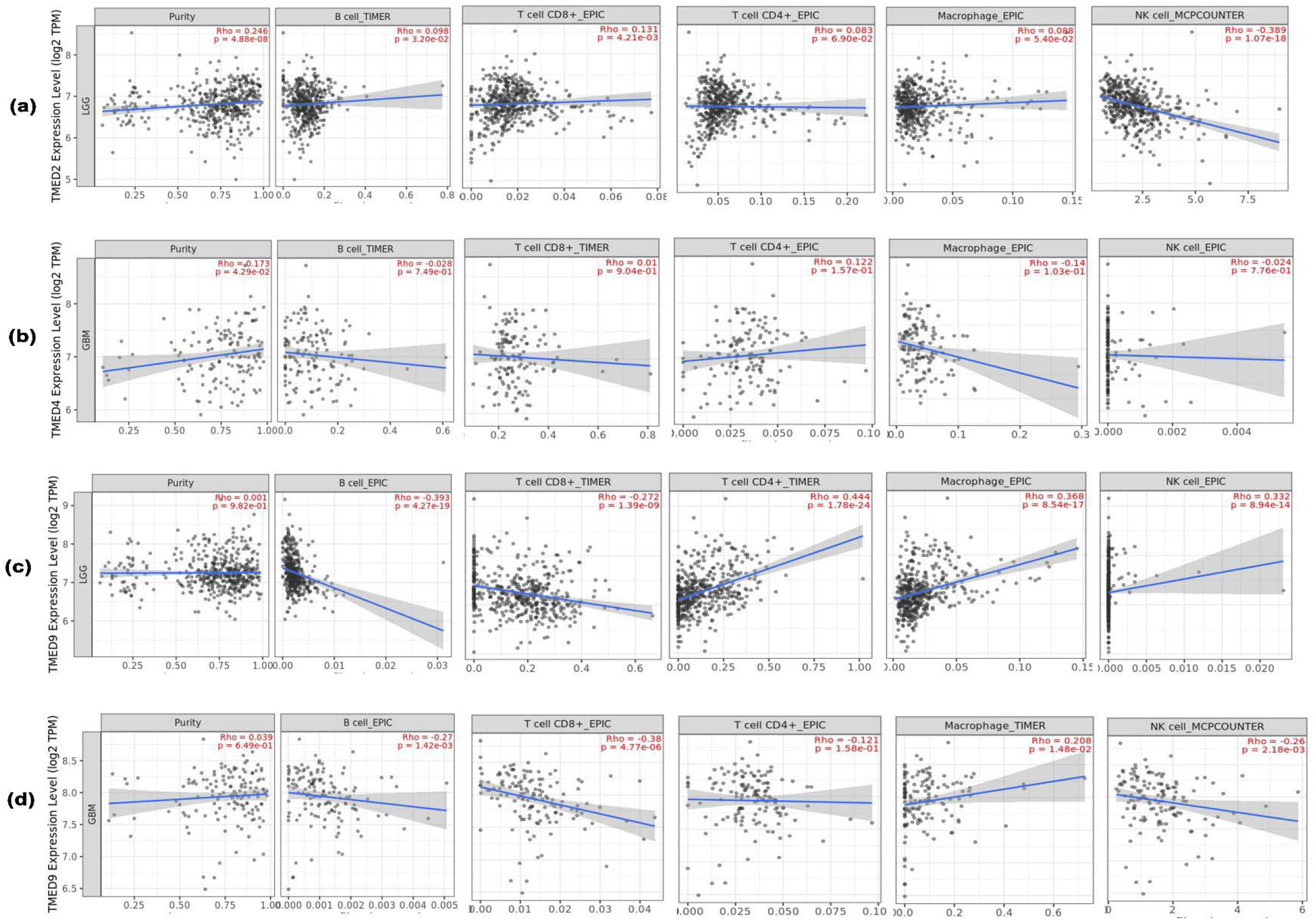
The association between *TMED4*and *TMED9* expression and immune cell infiltration levels in LGG (a) and GBM (b) patients. *TMED4*and *TMED9* expression was negatively and positively correlated with different immune cell abundance in glioma patients.

**Figure 8:**
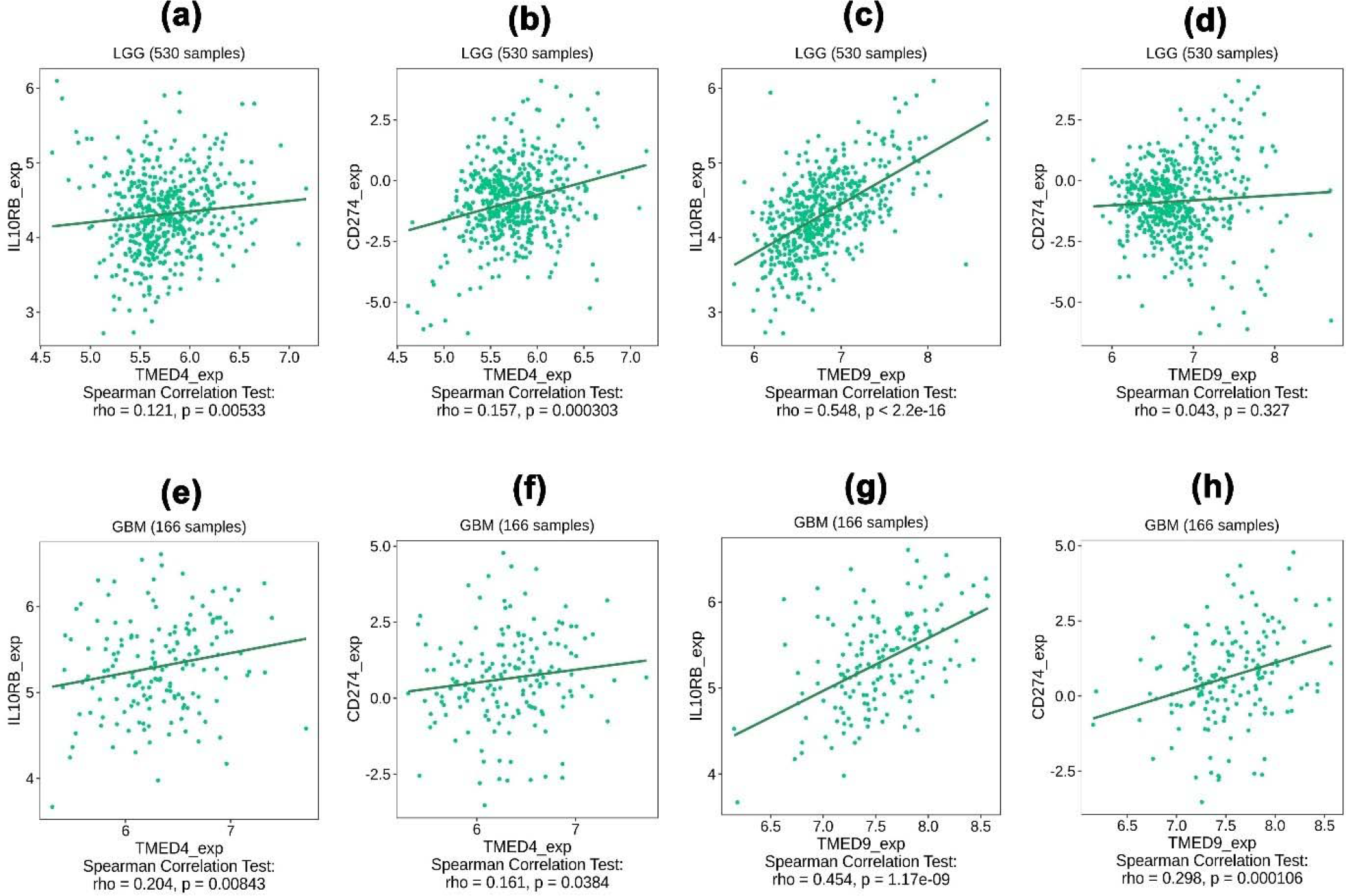
Association between the expression of the selected genes with different immunoinhibitors in LGG tissues: (a) *TMED4* vs IL10RB, (b) *TMED4* vs CD274, (c) *TMED9* vs *IL10RB*, (d) *TMED9* vs CD274 and in GBM tissues: *TMED4* vs IL10RB, (b) *TMED4* vs CD274, (c) *TMED9* vs *IL10RB*, (d) *TMED9* vs CD274.

### 3.7. The Positively Co-expressed Neighbor Genes of *TMED4* and *TMED9* in Glioma Patients and Their Functional Enrichment Analysis

The neighbor genes of *TMED4* and *TMED9* in LGG and GBM patients were identified using the LinkedOmicsserver (**Supplementary Figure S4**). *TMED4* was found to be highly and positively co-expressed with TRNA-YW Synthesizing Protein 1 Homolog (*TYW1*) (Cor: 0.73, p=1.84e-90) in LGG patients whereas it showed highest co-expression level with NudC Domain Containing 3 (*NUDCD3*) (Cor: 0.67, p=5.84e-21) genein GBM patients (**Figure 9**). On the other hand, *TMED9* showed highest co-expression level with Lectin, Mannose Binding 2 (*LMAN2*) both in the LGG (Cor: 0. 80, p=7.41e-117)and GBM (Cor: 0.71, p=1.73e-24)patients (**Figure 9**). Thereafter, the top 10 positively co-expressed genes of *TMED4* and *TMED9* in two different glioma tissues were selected for their functional enrichment analysis. The biological process analysis of the neighbor genes revealed that they were predominantly involved in collagen fibril organization, negative regulation of trophoblast and platelet degranulation (**Figure 10a**). Among the top selected molecular functions of the neighbor genes hexosyl transferase activity, transition metal ion binding and zinc ion binding were most significant (**Figure 10b**). The cellular component analysis revealed that the gene cluster was mostly operating in the focal adhesion point, cell-substrate junction and intracellular organelle lumen (**Figure 10c**). Different pathway analysis on the neighbor genes revealed that, most of them are involved in post-translational modification of different proteins, modulating protein functions in endoplasmic reticulum (ER), extracellular matrix organization, cell communication and so on (**Figure 10d, 10e** and**10f**).

**Figure 9:**
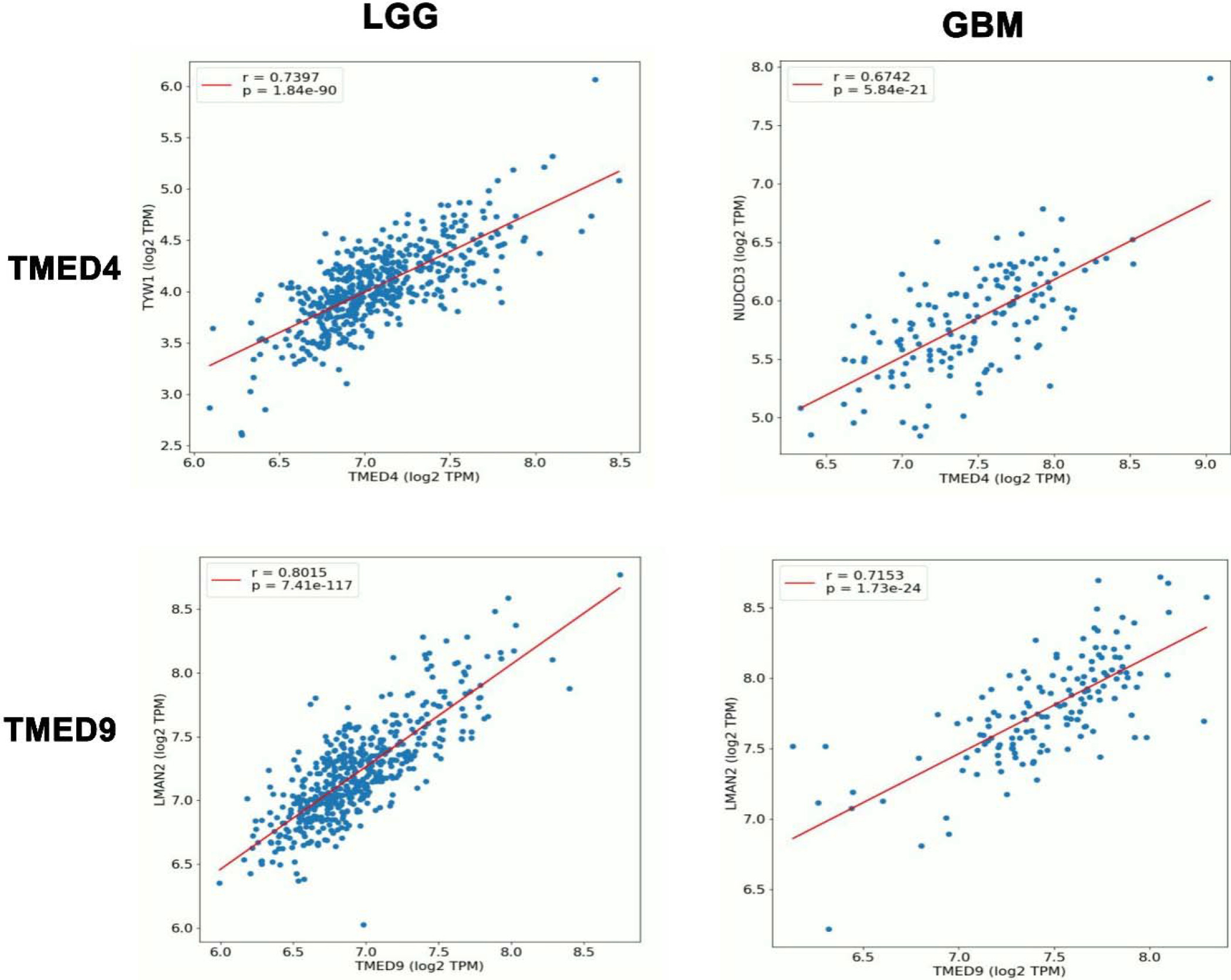
The top positively co-expressed genes of *TMED4* and *TMED9* in LGG and GBM tissues. *TMED4* showed co-expression with two different genes in LGG and GBM tissues whereas *TDME9* is highly co-expressed with one single gene in both LGG and GBM tissues.

**Figure 10:**
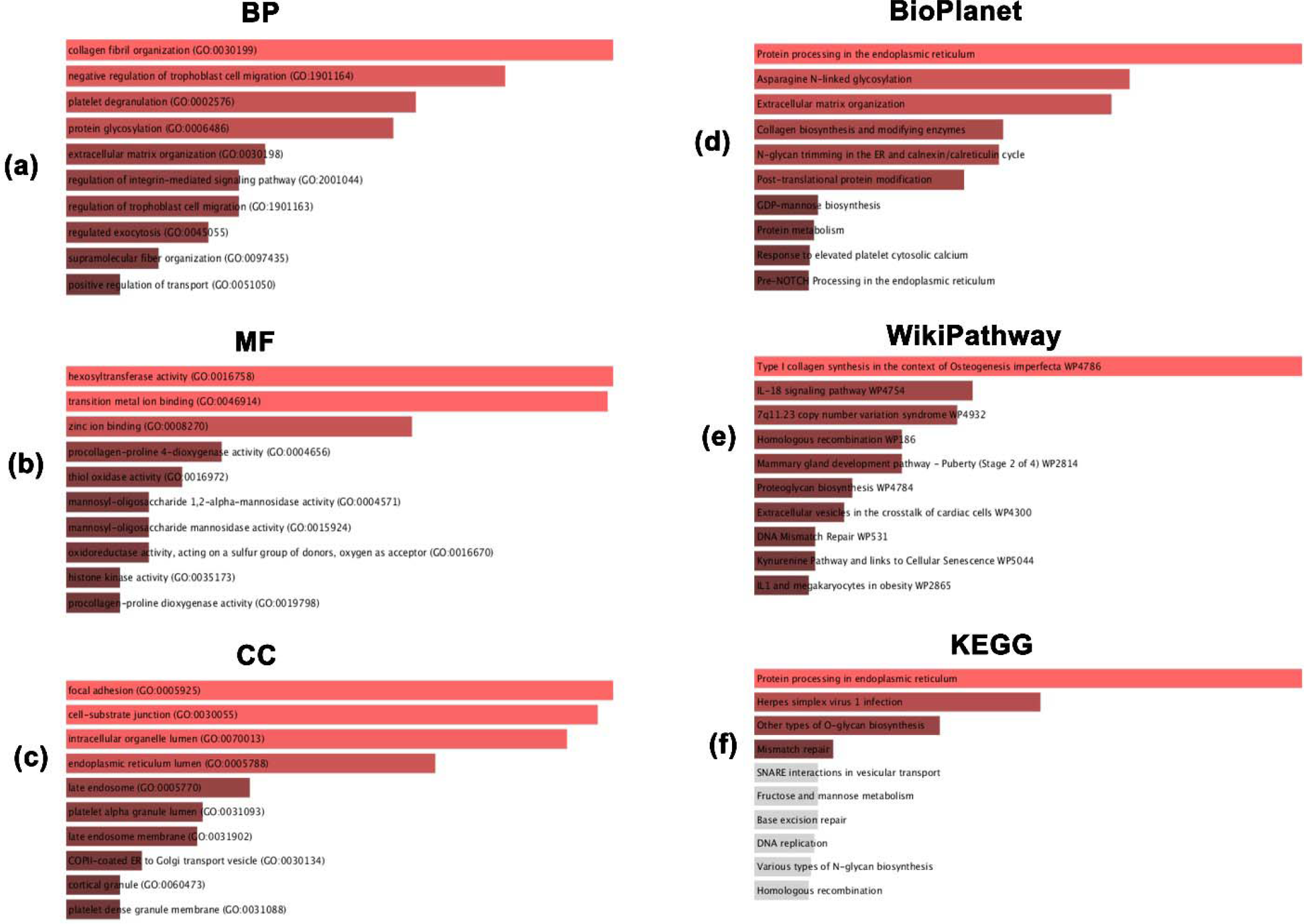
The top gene ontology terms i.e., BP: biological process (a), MF: molecular fuction (b) and CC: cellular component (c) of the co-expressed neighbor genes of *TMED4* and *TMED9* in LGG and GBM tissues. BioPlanet (d), WikiPathwa (e) and KEGG pathway analysis report on the top co-expressed neighbor genes of *TMED4* and *TMED9*. Red and dark-brown indicates significant association (p<=0.05) whereas ash color represents insignificant association (p>0.05).

## 4. Discussion

In this study, a database mining approach was utilized to evaluate and establish the prognostic value of *TMED4* and *TMED9* gene expression in glioma patients. Differential gene expression analysis can aid in understanding the roles of specific gene overexpression or under-expression in the oncogenic development of a cell and its subsequent growth [45]. Initially, *TMED4* and *TMED9* genes were found to be differentially expressed (upregulated) in LGG and GBM tissues at the mRNA and protein levels suggesting their possible oncogenic functions in glioma growth and progression (**Figure 1** and **2**). Nevertheless, they showed higher level of expression across different glioma cell lines. Since cancer development is a very complicated and multistep process varying across different cancer subtype, grade and patients’ demographic conditions [46], a comprehensive understanding on the expression level of a particular gene across different variables is required to understand its role in the oncogenic processes. In this study, our genes of interest showed increment in their expression with advancing age of glioma patients, glioma grades and histological subtypes which indicates that their expression might have significant association with glioma exacerbation and poor prognosis (**Figure 3)**.

DNA methylation is one of the major epigenetic drivers in cancer development and growth. Specifically, promoter methylation controls the gene activity of a particular gene by silencing or activating the transcription of that gene and hypomethylation is associated with upregulation of gene activity [47]. Coding sequence methylation also regulates the gene expression by altering the genome sequence structure in the chromatin architecture [48]. In this study, both the *TMED4* and *TMED9* promoter regions were found to be less methylated compared to the coding sequence regions, might be accounting for their overexpression in glioma tissues (**Figure 4)**. Moreover, the differential DNA methylation pattern of specific genes can serve in formulating methylation-sensitive diagnosis of glioma cells and making epigenetic clinical decision [49]. We also found that *TMED4* showed slight difference in methylation pattern between LGG and GBM tissues, whereas, *TMED9* maintains its distinct methylation feature even after transition from LGG to GBM state (**Figure 4)**. Therefore, *TMED9* may aid in the methylation-based diagnosis of glioma patients, whereas, *TMED4* is supposed to offer further stratification of LGG and GBM patients. Additionally, the association between *TMED9* hypomethylation and GBM patients’ poor OS and PFS signifies that it could also aid in the discovery of *TMED9*-based epigenetic glioma treatment measures (**Figure 4)**. However, such assumptions require further laboratory investigations.

Somatic driver mutations are the predominant causes of tumorigenic transformation of healthy cells inside human body [50,51], and somatic CNA contributes to a larger fraction in the cancer development than any other type of mutations [52]. These genetic changes influence the cancer initiation process by regulating oncogenes or tumor-suppressor genes [53]. Moreover, heterozygous amplification and deletion in specific genes like *EGFR* and *MDM2* can also serve as the molecular diagnostic target for high-throughput heterozygosity mapping in glioma patients [54,55]. Both *TMED4* and *TMED9* genes in glioma tissues were discovered to undergo multiple heterozygous CNA events with more amplification incidents in *TMED4* and deletion in *TMED9* which supports their potentiality as effective diagnostic candidates for glioma patients (**Figure 5)**. Moreover, the association between *TMED9* expression and different poor survival rates of glioma patients indicates that the CNA present in those genes might have critical underlying mechanisms in glioma development and prognosis that requires further investigation.

Thereafter, the survival analysis revealed that the higher expression of both the genes was negatively correlated with the OS of both LGG and GBM patients (**Figure 6)**. This suggests that the overexpression of *TMED4* and *TMED9* might have negative effects on glioma patients throughout different disease stages. This along with their higher expression in glioma tissues irrespective of patients’ demographic and clinicopathological conditions emphasizes that both the genes should aid in TMED-based tracking of glioma patients throughout the clinical courses.

Different immune cells along with adjacent intrinsic cell of the CNS forms the tumor microenvironment in gliomas which determines the cancer growth, metastasis and response to therapy [56,57]. Thus, different immune cells are prominent measures for formulating immunotherapy against glioma considering their ability to cross the blood brain barrier [58]. Different cytokine and chemokine secreting immune cells after getting produced in higher titers in the glioma microenvironment control the immunity against the glioma cells [59]. Abundance of numerous immunoinhibitors along with immune cells can assist in the diagnosis of glioma patients and keeping track on the patients throughout the disease state. For example, high abundance of CD8+ T cells in glioma patients predicts better survival [60]. In this study, *TMED4* and *TMED9* expression was found to be significantly correlated to the abundance of different immune cells (i.e., B cells and T cells) in LGG and GBM tissues that may aid in administering combinatorial immunotherapy along with TMED-based glioma treatment method or help in dual diagnosis (**Figure 7)**. The expression level of our genes of interest also showed significant association with the expression level of different immunoinhibitors i.e., PDC274, IL10RB (**Figure 8)**. Previously, CD274 infiltration level in glioma cells has been proposed to predict the favorable survival of glioma patients [61], as well as, IL10RB upregulation has been linked to the unfavorable survival of glioma patients [62].

The co-expression analysis revealed that *TMED4* gene is highly co-expressed with *TYW1* gene in LGG tissues and *NUDCD3* gene in GBM tissues (**Figure 9**). *TYW1* overexpression and mutations are associated with leukemia and breast cancer prognosis [63,64]. *NUDCD3* overexpression is also linked to the poor prognosis of cervical cancer patients [65]. *TMED9* gene was found to be highly co-expressed with *LMAN2* gene both in LGG and GBM tissues (**Figure 9**). Recent laboratory research has grounded that this particular gene is differentially methylated in brain white matter tissues and it is overexpressed in brain metastatic breast cancer tissues [66,67].

Pathway analysis of the co-expressed neighbor genes in glioma tissues indicated that most of the genes are involved in metal ion such as Zinc binding (**Figure 10**). Previous study suggests that deficiency in Zinc concentration, imbalance and deregulation is associated with different cancers including childhood brain tumor development [68]. Maintaining focal adhesion was among the top selected molecular functions of neighbor genes and suppression of such activity reduces brain cancer growth [69,70]. Cellular component analysis on the neighbor genes of *TMED4* and *TMED9* revealed that they are predominantly involved in protein processing in ER and any unfolded protein response in ER can influence the tumorigenesis in GBM tissues [71]. These evidences suggest that the neighbor genes of *TMED4* and *TMED9* in LGG and GBM tissues can also be investigated in glioma diagnosis and therapeutic measure discovery.

Overall, the reports in this study obtained from differential expression analysis, methylation and mutation frequency of *TMED4* and *TMED9* genes in LGG and GBM tissues indicate that these genes might have underlying mechanism in the glioma development and progression. Their overexpression pattern and effects on survival pattern signifies that the selected genes could be potential diagnostic and therapeutic target for glioma diagnosis and prognosis. Finally, this study recommends thar *TMED4* and *TMED9* are potential prognostic and therapeutic targets for glioma. However, further laboratory research is warranted to validate the significance of this study which is currently underway.

## 5. Conclusion

In summary, this study demonstrated the differential expression and variation in the methylation pattern of the promoter and coding sequences of *TMED4* and *TMED9* in LGG and GBM tissues. A number alteration events were reported in the coding regions of those genes in glioma tissues. Significant association between *TMED4* and *TMED9* overexpression and glioma patients OS was observed. These evidences suggest that *TMED4* and *TMED9* could be potential target for glioma diagnosis and treatment. Moreover, expression levels of multiple immune cells and immunoinhibitors were found to be associated with *TMED4* and *TMED9* expression in LGG and GBM tissues that may aid in formulating TMED-based diagnostic method and therapeutic interventions. The functional enrichment analysis revealed that their co-expressed genes are involved in functions and deregulation in those activity can promote gliomageneis. Thus, the neighbor genes of *TMED4* and *TMED9* could also be investigated further while extending laboratory work on making TMED-based diagnostic and therapeutic measures for glioma patients.

## Author’s Contribution Statement

MU conceived and designed the study. MU and AM carried out the experiment. MU and AM illustrated the graphs and figures. MU, TT and MF wrote the initial draft. MU, TT, MF, UZ and MR edited and revised the paper. MR supervised the study. All authors approved the final manuscript for publication.

## Ethics Approval and Consent to Participate

Not Applicable

## Consent for Publication

Not Applicable

## Availability of Data and Material

All the data are provided within the manuscript and the supplementary material.

## Acknowledgement

Authors are thankful to the members of Bio-resources Technology and Industrial Biotechnology Laboratory, Jahangirnagar University, Dhaka, Bangladesh and Swift Integrity Computational Lab, Dhaka, Bangladesh for their supports during the preparation of the manuscript.

