## Supplementary Information for "Expression Analysis, Molecular Characterization and Prognostic Evaluation on *TMED4* and *TMED9* Gene Expression in Glioma"

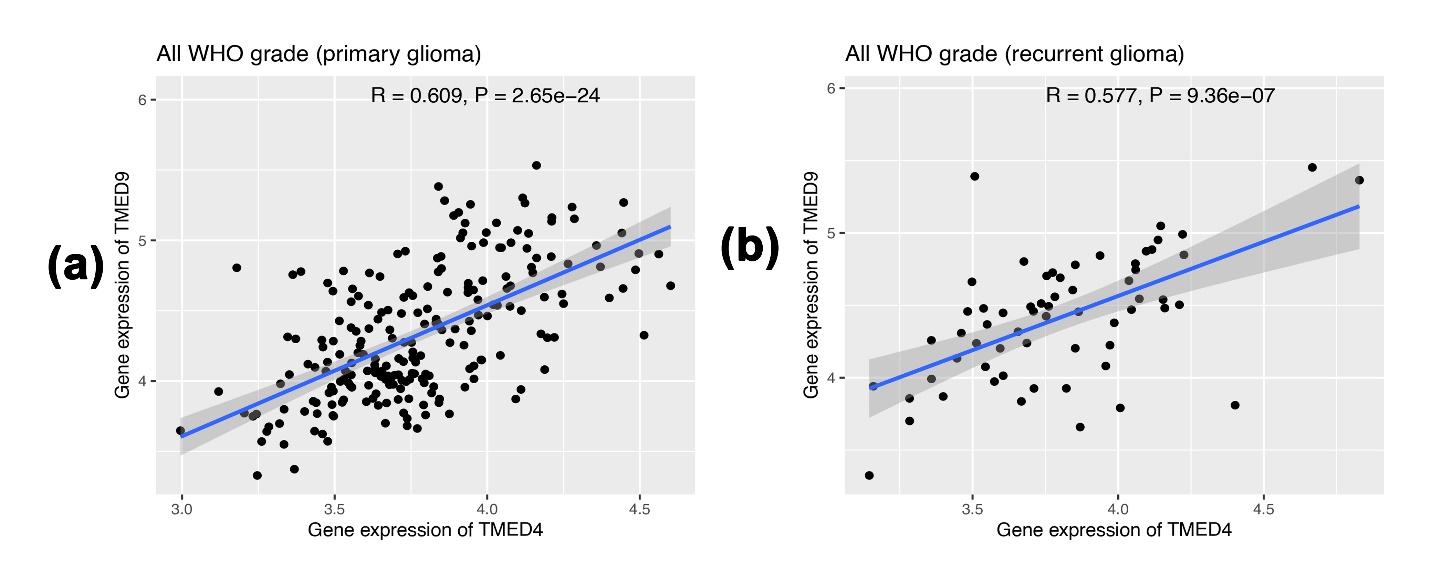
**Supplementary Information**

**Supplementary Figure S1:** The co-expression analysis of *TMED4* and *TMED9* genes in primary (a) and recurrent (b) glioma tissues.

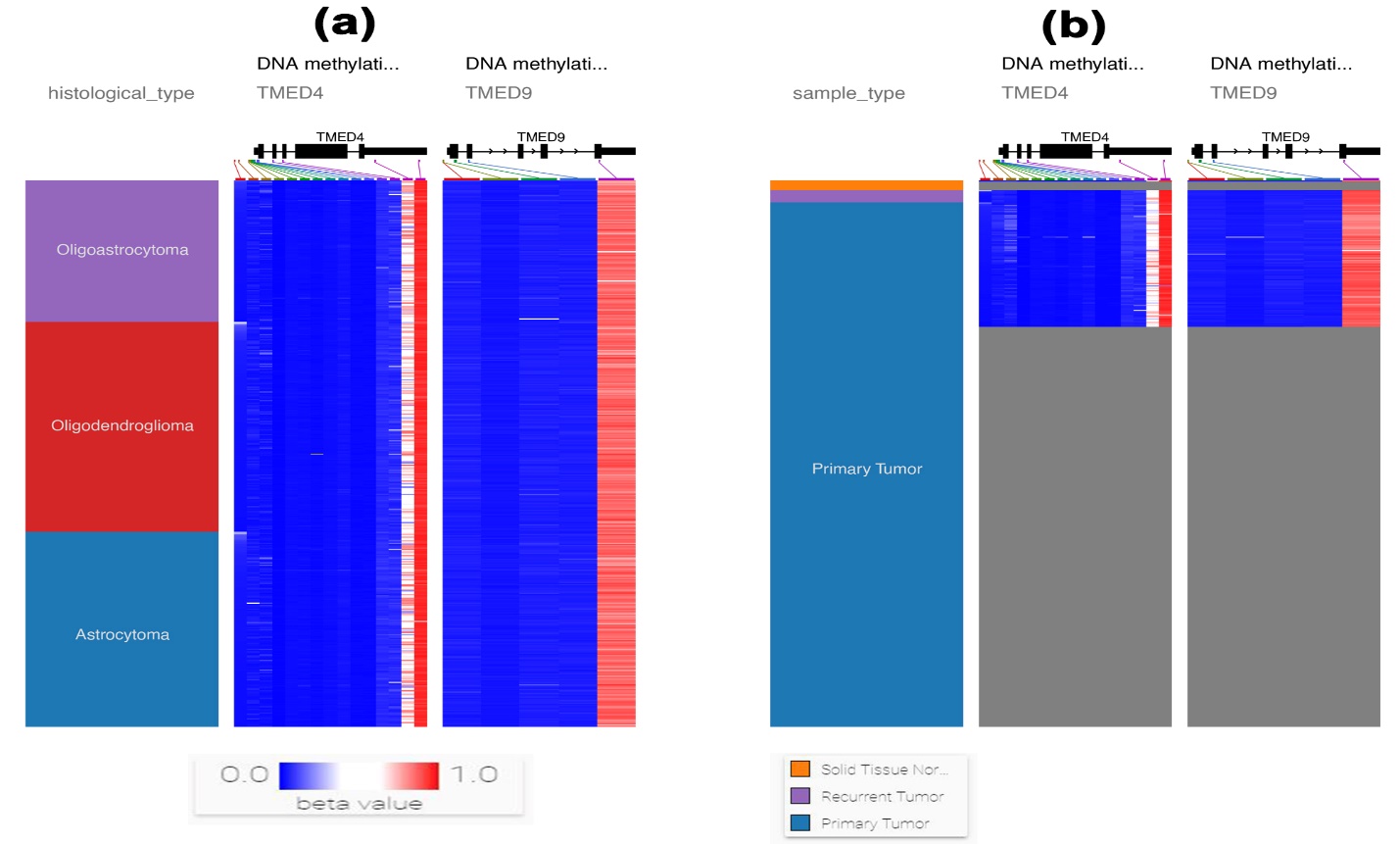

**Supplementary Figure S2:** The coding sequence methylation status of *TMED4* and *TMED9* genes in LGG (a) and GBM (b) tissues. Ash color indicates the missing of data corresponding to the groups presented in left column.

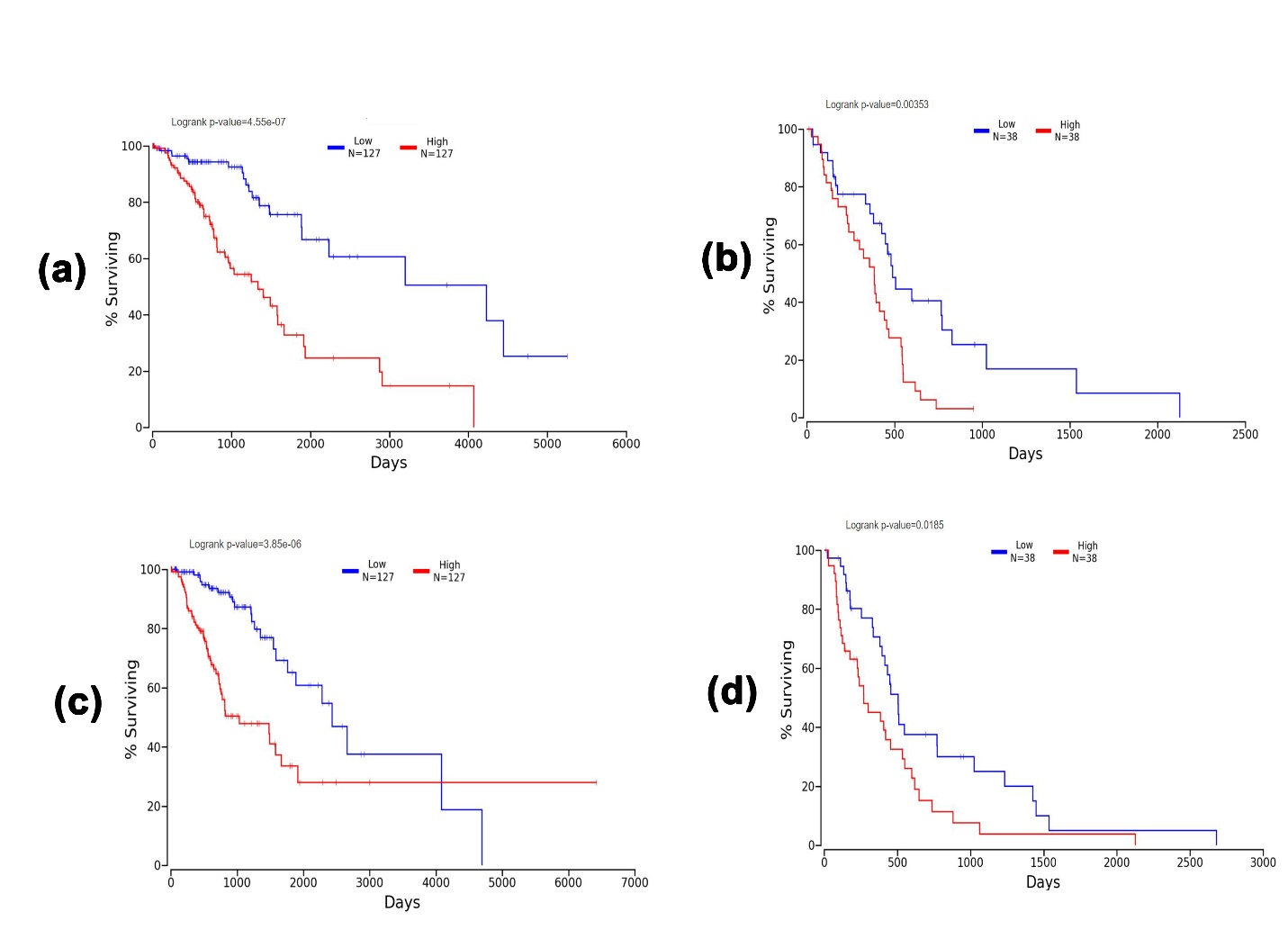

**Supplementary Figure S3:** The result of overall survival analysis on TMED4 gene expression in LGG(a) and GBM (b) patients from OncoLnc server. TMED4 overexpression was found to be negatively correlated with poor OS of LGG and GBM patients (p<0.05). The overall survival analysis on TMED9 gene expression in LGG (c) and GBM (d) patients which revealed negative correlation.

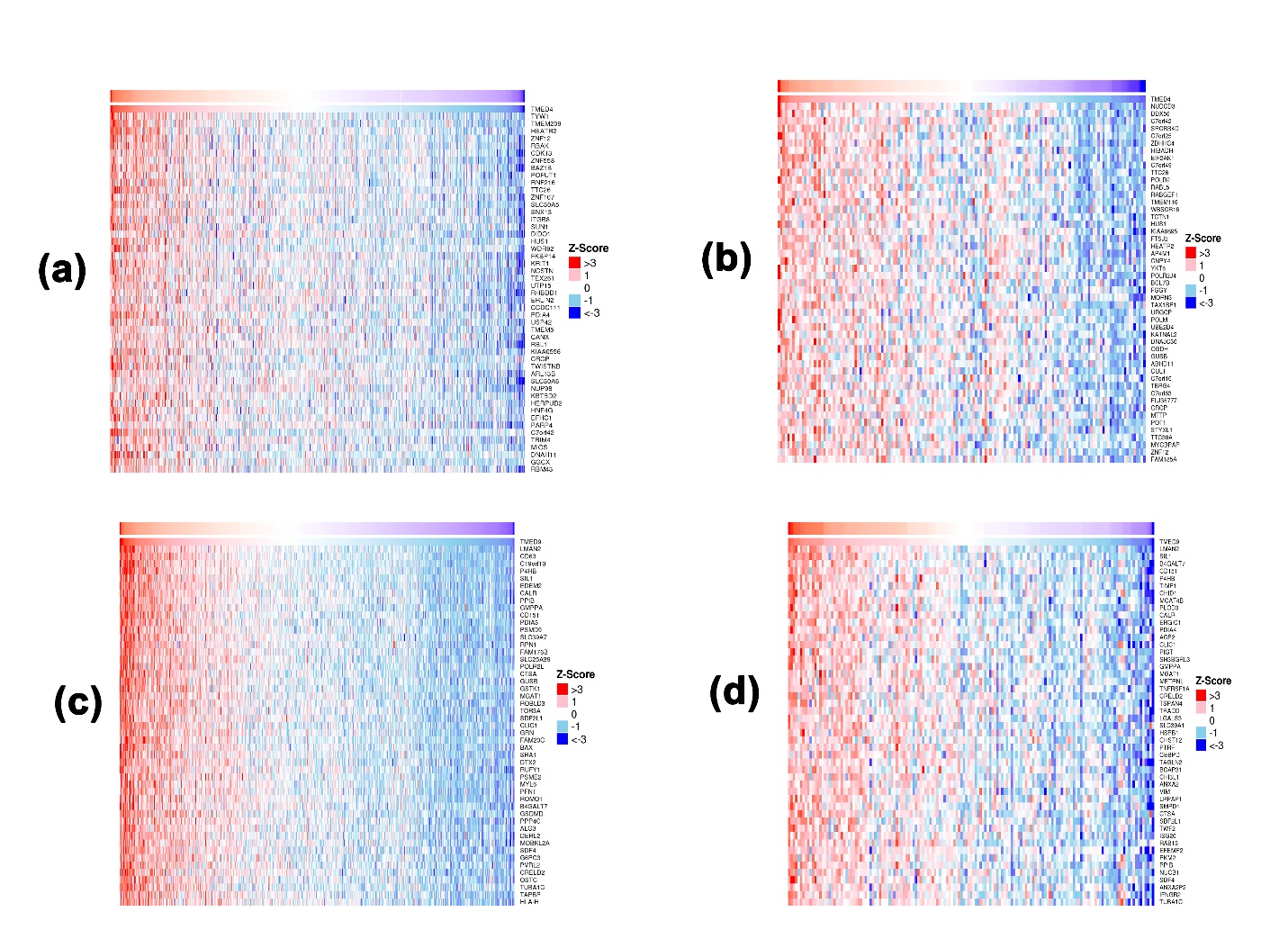

**Supplementary Figure S4:** The top co-expressed genes of *TMED4* in LGG (a) and GBM (b) tissues. The top co-expressed genes of *TMED9* in LGG (a) and GBM (b) tissues.

| **Cancer Type** | **Survival Type** | **P Value** |
| --- | --- | --- |
| *TMED4* | | |
| GBM | OS | 0.000654 |
|  | PFS | 0.000355 |
|  | DSS | 2.52E-05 |
| *TMED9* | | |
| GBM | OS | 0.318285 |
|  | PFS | 0.437907 |
|  | DSS | 0.448692 |
| *TMED4* | | |
| LGG | OS | 2.37E-07 |
|  | PFS | 4.02E-08 |
|  | DSS | 5.91E-08 |
|  | DFI | 0.28424 |
| *TMED9* | | |
| LGG | OS | 0.300927 |
|  | PFS | 0.831495 |
|  | DSS | 0.28185 |
|  | DFI | 0.87719 |

**Supplementary Table S1:** Summary of the survival analysis on glioma patients in relation to *TMED4* and *TMED9* mutation. OS: Overall Survival, PFS: Progression-free Survival, DSS: Disease-specific Survival, DFI: Disease-free Interval
